# ACE2-containing defensosomes serve as decoys to inhibit SARS-CoV-2 infection

**DOI:** 10.1101/2021.12.17.473223

**Authors:** Krystal L. Ching, Maren de Vries, Juan Gago, Kristen Dancel-Manning, Joseph Sall, William J. Rice, Clea Barnett, Feng-Xia Liang, Lorna E. Thorpe, Bo Shopsin, Leopoldo N. Segal, Meike Dittmann, Victor J. Torres, Ken Cadwell

## Abstract

Extracellular vesicles of endosomal origin, exosomes, mediate intercellular communication by transporting substrates with a variety of functions related to tissue homeostasis and disease. Their diagnostic and therapeutic potential has been recognized for diseases such as cancer in which signaling defects are prominent. However, it is unclear to what extent exosomes and their cargo inform the progression of infectious diseases. We recently defined a subset of exosomes termed defensosomes that are mobilized during bacterial infection in a manner dependent on autophagy proteins. Through incorporating protein receptors on their surface, defensosomes mediated host defense by binding and inhibiting pore-forming toxins secreted by bacterial pathogens. Given this capacity to serve as decoys that interfere with surface protein interactions, we investigated the role of defensosomes during infection by severe acute respiratory syndrome coronavirus 2 (SARS-CoV-2), the etiological agent of COVID-19. Consistent with a protective function, exosomes containing high levels of the viral receptor ACE2 in bronchioalveolar lavage fluid from critically ill COVID-19 patients was associated with reduced ICU and hospitalization times. We found ACE2+ exosomes were induced by SARS-CoV-2 infection and activation of viral sensors in cell culture, which required the autophagy protein ATG16L1, defining these as defensosomes. We further demonstrate that ACE2+ defensosomes directly bind and block viral entry. These findings suggest that defensosomes may contribute to the antiviral response against SARS-CoV-2 and expand our knowledge on the regulation and effects of extracellular vesicles during infection.

## INTRODUCTION

Exosomes are a subgroup of single membraned vesicles 40-120 nm in diameter secreted by virtually every cell type. Cargo molecules ranging from non-coding RNAs to proteins involved in signal transduction are incorporated during exosome biogenesis, which involves exocytosis of endosomal structures [1]. As such, exosomes mediate intercellular communication events involved in cell proliferation, migration, and cancer [2]. A role in host defense is supported by the finding that exosomes can deliver nucleic acids and other immunogenic moieties from infected cells that elicit interferon responses or promote antigen presentation by target cells [3-8]. However, how exosomes are regulated in response to infections remains unclear. A better understanding may inform strategies that seek to use extracellular vesicles as biomarkers or therapeutic agents for infectious disease.

In our previous work, we identified exosomes involved in host defense termed defensosomes that mediated protection against bacterial pore-forming toxins that kill target cells, such as α-toxin produced by *Staphylococcus aureus* [9]. Recognition of bacterial DNA by toll-like receptor 9 (TLR9) led to increased production of defensosomes decorated by ADAM10, the host cell surface target of α-toxin, in a manner dependent on ATG16L1 and other components of the membrane trafficking pathway of autophagy. We further demonstrated that ADAM10^+^ defensosomes serve as decoys that bind α-toxin and prevent cytotoxicity in cell culture and animal models [9]. Therefore, in addition to mediating intercellular communication by transferring signaling molecules between cells, exosomes can also promote host defense through a surface interaction with bacterial proteins. Whether defensosomes are deployed during a viral infection remains unclear.

Severe acute respiratory syndrome coronavirus 2 (SARS-CoV-2), the respiratory virus responsible for the ongoing global COVID-19 pandemic, can cause life threatening tissue injury to the lung as well as extrapulmonary symptoms that are less understood [10]. Entry into host cells mainly depends on the binding of receptor-binding domain (RBD) of SARS-CoV-2 spike (S) protein to host receptor ACE2. The affinity of SARS-CoV-2 spike for ACE2 is 5 times higher than that of SARS-CoV spike [11], and 1,000 times higher than hemagglutinin of Influenza A for sialic acid [12]. Given this dependence on a high affinity cell surface interaction for viral entry, we hypothesized ACE2 containing defensosomes would inhibit SARS-CoV-2 infection through the binding of the spike protein.

## RESULTS

### Increased proportion of ACE2+ exosomes in human BAL fluid correlates with reduced time in the ICU

Few studies, if any, have analyzed exosomes in mucosal tissues of patients during an ongoing infection. This is especially true in the setting of SARS-CoV-2 infection where access to lower airway samples were limited due to concerns about possible risks for health care professionals [13-20]. In the airways, *ACE2* expression exists as a gradient, with the highest levels in the nasal passageway, down to the lower airways where it is exclusively expressed on Type II pneumocytes [21]. To determine whether ACE2+ exosomes are generated and are present in the respiratory tract, we analyzed bronchioalveolar lavage fluid (BALF) samples from 80 critically ill COVID-19 patients [22] using biochemical fractionation to enrich for exosomes followed by flow cytometry [9]. All patients in our cohort were admitted to the intensive care unit (ICU) due to respiratory failure requiring invasive mechanical ventilation and tested positive for SARS-CoV-2 infection [22]. Applying stringent gating criteria for exosomes, we observed remarkable interindividual variation in the proportion of exosomes with surface ACE2 and the levels of ACE2 present on each exosome measured by mean fluorescence intensity (MFI) (Fig. 1A, Supp. Fig. 1A-C). We found that 10 patients had at least two-fold higher exosomes levels in their BALF compared to the mean exosome levels from the other patients. These data suggest that there is variation between individuals in how much ACE2 is loaded into each exosome as well as how many ACE2+ exosomes are produced by each person.

**Figure 1.**
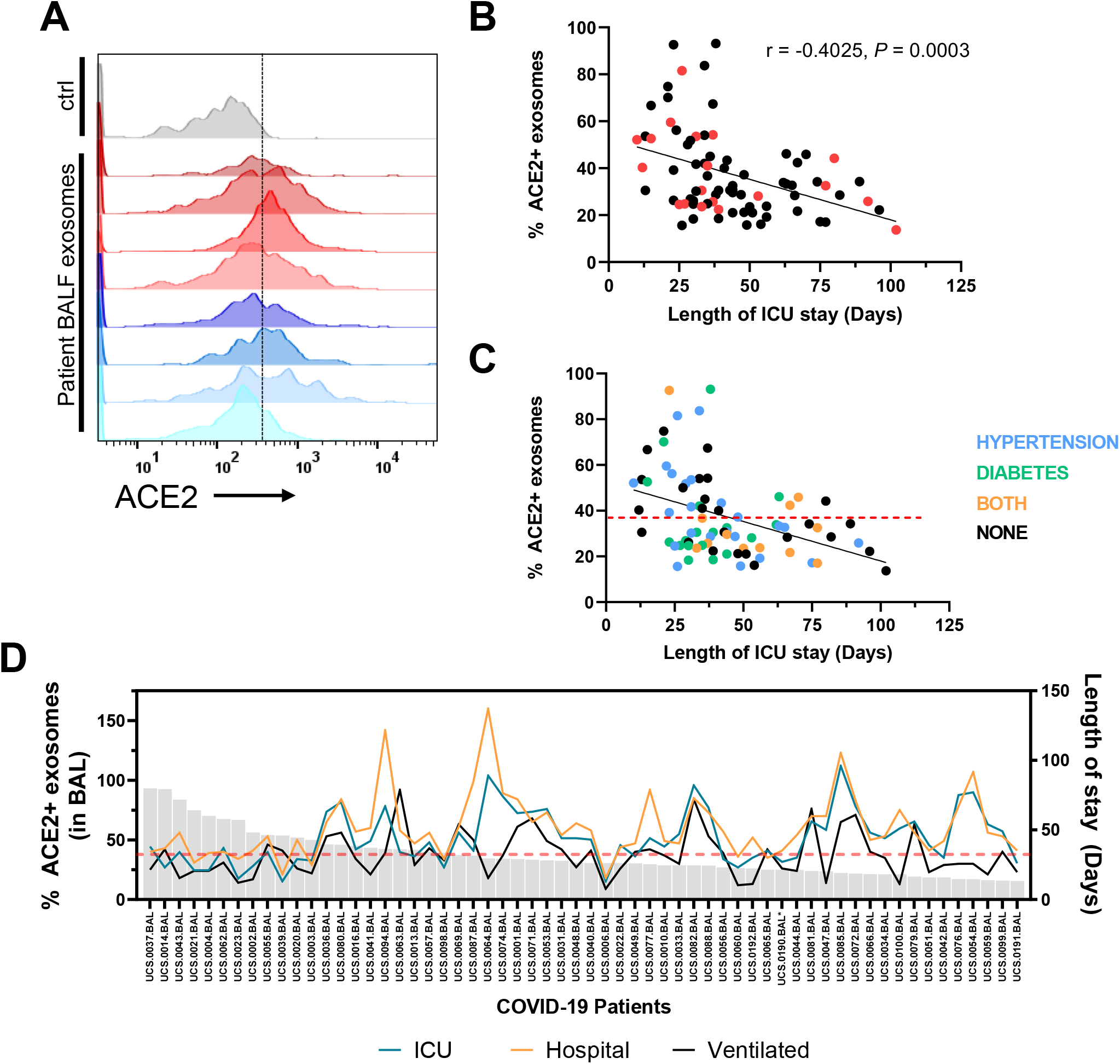
ACE2+ exosomes are associated with reduced length of stay in ICU for COVID-19 patients. **(A)** Flow cytometry histograms of ACE2 levels on exosomes in BALF from 8 representative COVID-19 patients. Ctrl: isotype control on exosomes from non-covid patient BALF. **(B)** Correlation analysis of proportion of ACE2+ exosomes in BALF and length of stay in the ICU *(N=78)*. Red dots indicate deaths. Simple regression analysis. **(C)** Correlation analysis of % ACE2 exosome levels with comorbidities colored: Hypertension (blue), Diabetes (green), both (orange). Red line indicates the average % of ACE2+ exosomes of all patients. **(D)** Length of stay in the ICU (blue), total hospitalization time (orange), and time on a ventilator (black) for each COVID patient (excluding deaths) plotted against the proportion of ACE2^+^ exosomes isolated from BALF *(N=59)*. r, Pearson correlation coefficient.

We then asked whether level of ACE2 on exosomes or proportion of ACE2+ exosomes correlate with clinical parameters such as patient age, sex, disease severity, infection status, and other comorbidities linked to SARS-CoV-2 infection such as diabetes or hypertension. 76.2% of the patients were male, mostly nonwhite (53.8%), with a mean age of 62 years (SD, 14.2). Hypertension and diabetes were reported in 52.5% and 38.3% of patients, respectively (Supp. Table 1). Two patients were excluded from analysis because they were not within 3 standard deviations from the mean (47.96 days) for length of stay in the ICU. As a sensitivity analysis, we also ran the regressions using ACE2 MFI as a predictor of ICU stay outcome, resulting in the same trends in the covariates (Supp. Table 3). Compiling the information regarding the proportion of exosomes that contain ACE2+ in patient BALF, we ran different models using clinical data to test whether there was any association with ACE2+ exosomes and length of stay in the ICU, as there was no correlation with mortality. We performed a linear model and a negative binomial model to test this relationship, and we controlled for age, gender, culture results of BAL and blood, diabetes, and hypertension. We also included in a sensitivity analysis model an interaction term between gender, sex, and hypertension, as they are usually correlated. We expressed the coefficients without interactions for simplicity because the estimates did not significantly change. The linear model resulted in negative correlation between length of stay in the ICU and the proportion of ACE2+ exosomes found in the BALF (Table 1, Fig. 1B). The negative binomial model predicted similar results (Supp. Table 2). To account for increased length of stay in the ICU due to complications or subsequent infections, we also used ventilation days as an outcome, and found that proportion of ACE2+ exosomes remained a significant predictor (Supp. Tables 4-5).

**Table 1.** Predictors of length of stay in the ICU among COVID-19 patients using a linear regression model.

| Length of stay in ICU (Days) |  |  |  |
| --- | --- | --- | --- |
| Predictors | Estimates | CI | P Value |
| Age (years) | 0.2338 | -0.5709 – 1.0385 | 0.564 |
| % ACE2 positive (BAL exosomes) | -0.549 | -0.7845 – -0.3136 | <0.001 |
| Sex [M] | 39.7312 | -13.3973 – 92.8596 | 0.14 |
| Hypertension | 12.1199 | -81.5270 – 105.7668 | 0.797 |
| BAL *C. albicans* [Positive] | 21.05 | 6.7861 – 35.3138 | 0.004 |
| Blood culture final result [Positive] | 5.4402 | -3.8165 – 14.6968 | 0.245 |
| (Intercept) | 50.0676 | 2.4603 – 97.6750 | 0.04 |
| **Observations** | 78 |  |  |
| **R<sup>2</sup> / R<sup>2</sup> adjusted** | 0.375 / 0.281 |  |  |

These analyses uncovered a threshold where patients with a proportion of ACE2+ exosomes lower than the mean (38.78%) generally have increased lengths of stay in both the ICU and hospital, and increased days on a ventilator (Fig. 1D, Supp. Fig. 1D). Individuals with diabetes (or both diabetes and hypertension) clustered below the average proportion of ACE2+ exosomes (Fig. 1D), suggesting common comorbidities may influence the proportion of ACE2+ exosomes. Individuals with hypertension have been shown to have reduced levels of circulating ACE2 with the adverse consequence of increased concentration of the vasoconstricting molecule Angiotensin II due to activation of ACE [23]. However, neither diabetes nor hypertension were associated with statistically significant differences in ACE2 levels on exosomes (Supp. Fig. 1E-F). Altogether, these data show that the proportion of ACE2+ exosomes and level of ACE2 on individual exosomes in BALF from patients with COVID-19 are highly variable, and their increased presence are associated with a reduction in the hospitalization days.

### Defensosomes are produced in response to SARS-CoV-2 and antiviral sensors

We previously showed that exosomes are induced by TLR9 activation in response to bacterial DNA or synthetic agonist CpG-A. It was unclear whether innate immune ligands generated during RNA viral infections affect exosome production. Although SARS-CoV-2 is not predicted to directly activate DNA sensors, damage during respiratory tract infection can lead to production of oxidized mitochondrial DNA (ox-mtDNA) [24, 25], which is known to be a potent ligand for TLR9 [26]. Stimulation with ox-mtDNA increased exosome production in A549 cells, but this was not observed with sheared genomic DNA (gDNA) or DNA fragments without any oxidized bases (Fig. 2A). We also tested CpG-B, which can activate TLR9, but unlike CpG-A, lacks a phosphodiester backbone and poorly induces interferons (IFNs) [27]. CpG-B stimulation failed to elicit exosomes from A549 cells (Fig. 2B). SARS-CoV-2 has been shown to indirectly activate the cytosolic DNA sensing pathway dependent on cGAS/STING [28]. However, ligands for cGAS/STING (2’3’-cGAMP and 3’3’-cGAMP) failed to induce exosomes (Fig. 2B).

**Figure 2.**
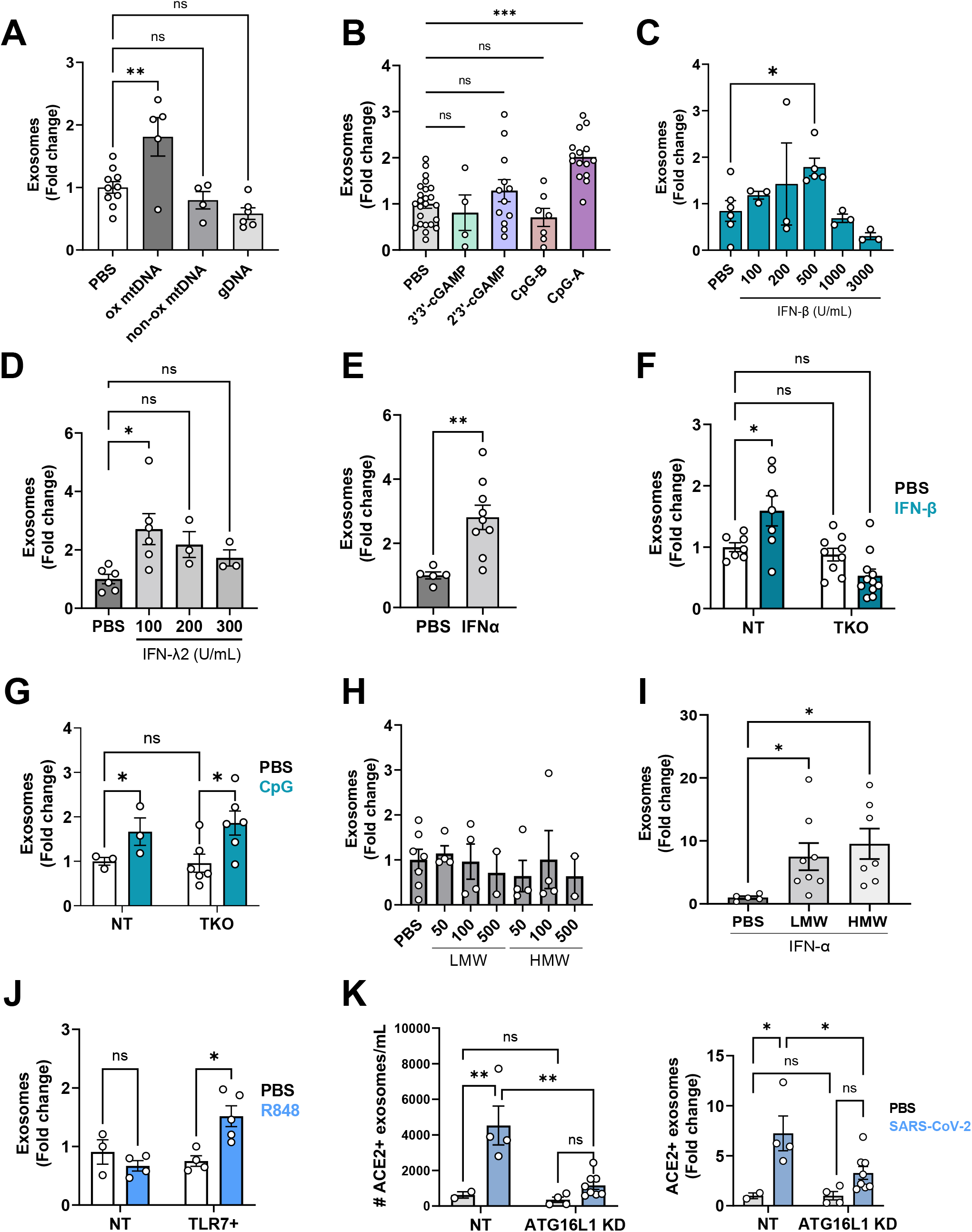
TLR ligands and SARS-CoV-2 induce exosome production. **(A)** Quantification of exosomes from A549 cell-culture supernatant by flow cytometry 16 hr after stimulation with 32 μg of oxidized mtDNA (ox mtDNA), mtDNA products without oxidized bases (non-ox mtDNA) or genomic DNA (gDNA) (*n=4-10*). **(B)** Quantification of exosomes from A549 cells stimulated for 16hr with 1μg/mL 3’3’-cGAMP *(n=4)*, 2’3’-cGAMP *(n=9)*, 0.5μM CpG-A *(n=8)*, and 0.5μM CpG-B *(n=7)*. CpG-A will be referred to as CpG from this point on. **(C)** Quantification of exosomes from A549 cells stimulated with various dilutions of IFN-β for 16hr (*n=3-6*). **(D)** Quantification of exosomes from A549 cells stimulated with various dilutions of IFN-*λ* for 16hr (*n=3-6)*. **(E)** Quantification of exosomes following stimulation of A549 cells with 100U/mL IFNα for 16hr *(n=5-9)*. **(F)** Exosome quantification of IFNAR/γR/λR triple KO (TKO) A549 cells stimulated with 500 U/mL IFN-β for 16hr compared to non-transduced (NT) A549 cells *(n=7-11)*. **(G)** Quantification of exosomes from nontransduced (NT) A549 or TKO A549 cells stimulated with CpG for 16hr *(n=3-6)*. **(H)** Quantification of exosomes from A549 cells stimulated with high or low molecular weight (HMW, LMW) Poly I:C with LyoVec at 50, 100, 500ng/mL *(n=2-7)*. **(I)** Exosomes were quantified from A549 cells pretreated with 100U/mL IFNα for 24hr, followed by stimulation with 10μg/mL LMW or HMW PolyI:C. **(J)** Quantification of exosomes from A549 cells and TLR7 expressing A549 cells stimulated with 1μg/mL R848 for 16 hr. (*n=3-5*). **(K)** ACE2^+^ A549 (NT) and ACE2^+^ A549 cells knocked down for ATG16L1 using shRNA (ATG16L1 KD) were infected with TCID50 SARS-CoV-2 USA-WA1/2020 for 72 hrs. Exosomes were isolated from supernatants and quantified in exact counts (left) and fold change (right). (**A-K)**: Exosomes: CD9+ CD63+ CD81+ events. Measurements were taken from distinct samples. Error bars represent mean ± SEM and at least 2 independent experiments were performed. **a**,**b**,**c**,**d**,**e**,**h**,**j** One-way ANOVA with multiple comparisons with Dunnett’s post-test. **f**,**g**,**i**,**k** Two-way ANOVA using Dunnett’s post-test. * *P* ≤ 0.05; ** *P* ≤ 0.01; ns: not significant.

It is possible that IFNs generated downstream of TLR9 activation mediate exosome production. Stimulation of A549 cells with IFN-β induced exosomes in a concentration dependent manner, where at higher concentrations, the number of exosomes began to decrease (Fig. 2C), which may be due to ISGylation and degradation of the compartment where exosomes are generated [29]. IFN-α and IFN-λ2 also induced exosomes (Fig. 2D-E). As expected, IFN-β failed to elicit exosomes from triple-knockout (TKO) A549 cells that lack receptors necessary for recognition of all three major classes of IFNs (IFNAR, IFNGR, and IFNLR) (Fig. 2F). In contrast, TKO A549 cells were still able to produce exosomes in response to CpG-A (Fig. 2G). Thus, IFN stimulation is sufficient to induce exosome production, but are not required downstream of nucleic acid sensing.

SARS-CoV-2 infection generate viral ssRNA and dsRNA that may serve as signals to induce exosomes. When transfected into cells, low and high molecular weight versions of the dsRNA mimetic poly I:C preferentially activate the cytosolic RNA sensors RIG-I and MDA-5, respectively, but we found that neither induced exosomes (Fig. 2H. In A549 cells, the endosomal dsRNA sensor TLR3 is expressed at low levels; IFN-α treatment induces TLR3 to functionally active levels, allowing A549 cells to become sensitive to untransfected poly I:C [30]. Following pretreatment with IFNα for 24hr, A549 cells produced 10-20-fold more exosomes in response to poly I:C without a transfection agent (Fig. 2I). A549 cells do not express the endosomal ssRNA sensors TLR7 or TLR8. We found that ectopic expression of TLR7 led to exosome induction upon stimulation with a synthetic agonist, R848 (Fig. 2J).

Given that multiple innate immune ligands induce exosome production, we asked whether ACE2+ exosomes are induced by SARS-CoV-2 infection. We found that SARS-CoV-2 infection of A549 cells ectopically expressing ACE2 (A549^ACE2+^) induced robust production of ACE2+ exosomes (Fig. 2K). Similar to our observations with bacteria, SARS-CoV-2-induced exosome production was impaired by *ATG16L1* knockdown (Fig. 2K). Therefore, the enhanced production of ACE2+ exosomes display a key hallmark of defensosomes – induced in response to infectious cues in a manner dependent on ATG16L1.

### ACE2+ defensosomes neutralize SARS-CoV-2 virions

Our finding that ACE2+ exosomes in patient BALF display an inverse correlation with length of hospital stay raised the possibility that they play a role during viral infection akin to our previous description of defensosomes during bacterial toxinosis. Therefore, we tested whether addition of exosomes can protect Vero E6 cells, which are highly susceptible to SARS-CoV-2 infection (Fig. 3A). Exosomes were isolated from A549^ACE2+^ cells (Fig. 3B) and mixed with human SARS-CoV-2 USA WA1/2020 isolate at an MOI of 0.01. We included a polyclonal neutralizing antibody targeting spike (nAb) as a positive control. Addition of ACE2+ exosomes led to a dose-dependent reduction in viral nucleoprotein (NP) staining by immunofluorescence microscopy at 24 hr post-infection (Fig. 3C-D, Supp. Fig. 2A), whereas ACE2-exosomes from untransduced control A549s failed to inhibit infection. Also, addition of the microvesicle fraction that does not contain exosomes (5000x*g*) did not decrease infection levels (Fig. 3C). We also tested neutralization in human airway epithelial cultures (HAECs), an air-liquid (ALI) interface model that contains all major cell types found in the bronchial epithelium (basal, ciliated, secretory), organized in pseudostratified architecture [31]. Addition of ACE2+ exosomes mixed with SARS-CoV-2 to the apical side (“lumen”) blocked infection of HAECs, while ACE2-exosomes did not show any significant effect (Fig. 3E, Supp. Fig. 2B).

**Figure 3.**
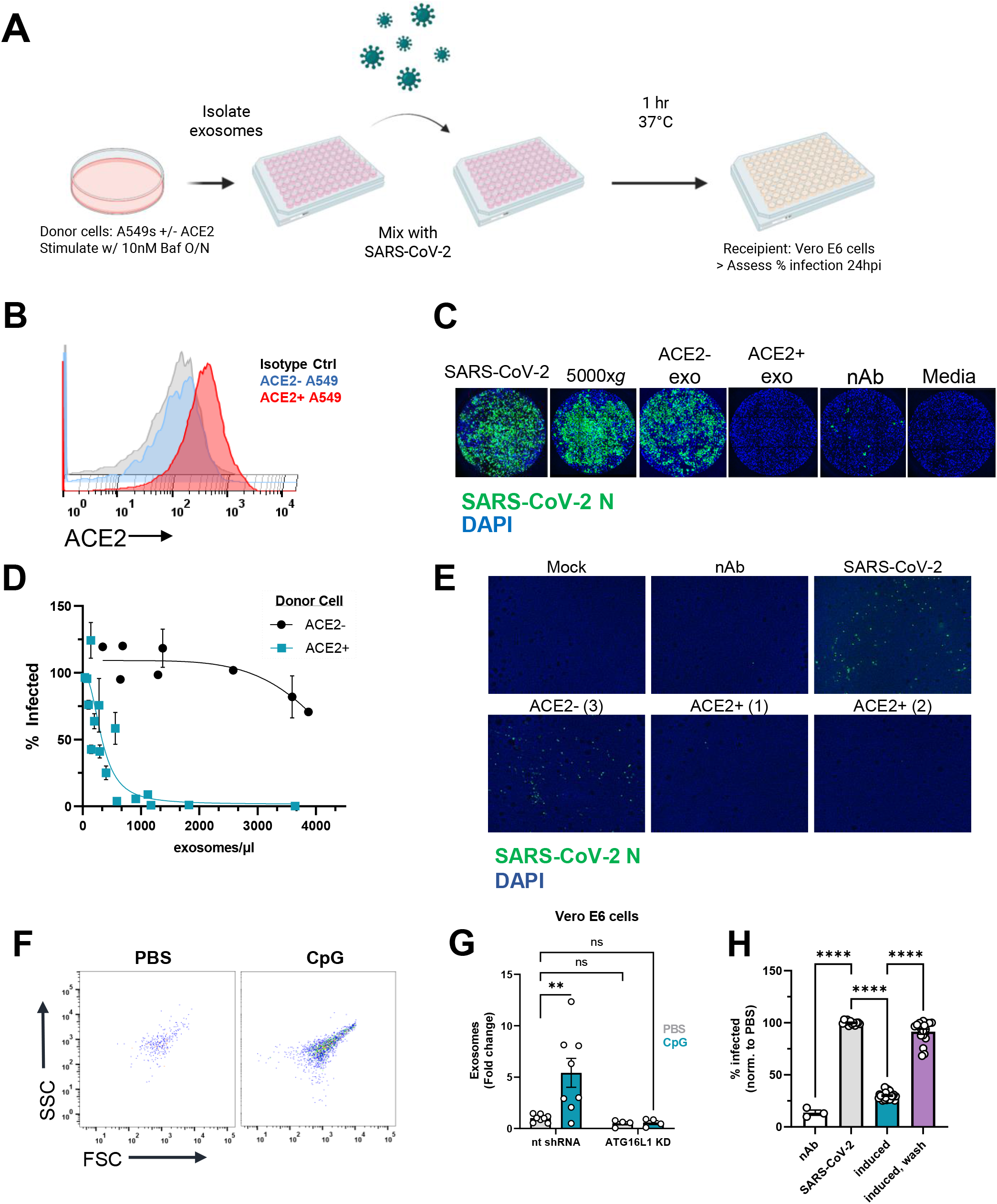
ACE2+ exosomes protect against SARS-CoV-2 infection. **(A)** Workflow for exosome neutralization assay used to assess protective effects of mixing SARS-CoV-2 and exosomes. Created with BioRender.com. **(B)** Representative histogram of cell-surface ACE2 on A549 and A549^ACE2^ cells. **(C)** Representative immunofluorescence images of Vero E6 cells infected with SARS-CoV-2, pellet from 5000x*g* spin, ∼20K exosomes from ACE2-cells, ∼20K exosomes from ACE2+ cells, neutralizing antibody, or with media alone stained for viral N protein. **(D)** Percentage of infected Vero E6 cells based on positive staining for N protein after infection with SARS-CoV-2 mixed with exosomes isolated from A549^ACE2^ (teal) or untransduced A549 cells (black). Compiled means ± SEM from 5 experiments, values normalized to infection with SARS-CoV-2 alone. (**E**) Representative immunofluorescence images of HAECs infected with mock (media only control), SARS-CoV-2 alone, ∼11K (1) or ∼22K (2) ACE2+ exosomes with virus, ∼150K (3) ACE2-exosomes with virus, and neutralizing antibody with virus (nAb). Images were taken 72h.p.i. **(F)** Representative flow cytometry plots of exosomes isolated from Vero E6 cells stimulated with CpG. FSC: forward scatter, SSC: side scatter. **(G)** Quantification of exosomes from Vero E6 cells and Vero E6 cells transduced with a non-targeting shRNA for ATG16L1 stimulated with PBS *(n=4)* or 1uM CpG-A *(n=4)* for 16 hr. **(H)** Quantification of infected cells after infection of Vero E6 cells with SARS-CoV-2 only, SARS-CoV-2 mixed with supernatants of cells pretreated overnight with CpG-A (induced), or SARS-CoV-2 added to CpG-A pretreated cells following removal of supernatant (wash). Exosomes: CD9, CD63, CD81+, PKH67+ events. Means ± SEM of at least 2 independent experiments. **g** Two-way ANOVA with Dunnett’s post-test compared to NT PBS. **h** One-way ANOVA with Dunnett’s post-test compared to PBS. Error bars represent the mean ± SEM of at least 2 independent experiments, with measurements taken from distinct samples. ** *P* ≤ 0.01; **** *P* ≤ 0.0001. ns: not significant.

We next tested whether inducing exosome production protects against viral infection without the need of adding exogenous exosomes from donor cells. We found that pre-treatment with CpG-A, which we confirmed induces exosome production by Vero E6 cells in a manner dependent on ATG16L1 (Fig. 3F-G), led to a reduction in cells infected with SARS-CoV-2 (Fig. 3H). Removing the exosome-containing supernatant prior to infection led to similar levels of SARS-CoV-2 infection as unstimulated cells (Fig. 3H, Supp. Fig. 2C). This protection is unlikely to be mediated by IFNs, as Vero E6 cells cannot produce IFNs [32, 33]. Together, these results show that defensosomes can inhibit SARS-CoV-2 infection.

### ACE2+ defensosomes directly interact with SARS-CoV-2

We next used electron microscopy techniques to visualize the interactions between SARS-CoV-2 virions and defensosomes. Viral particles with predicted spherical shape and uniform size of 80-100 nm in diameter were detected by negative stain imaging in control suspensions containing SARS-CoV-2 virions alone (Fig. 4A) [34]. With the addition of exosomes, we detected structures with the characteristic collapsed morphology of exosomes clustered around particles with virion morphology. The difficulty in distinguishing exosomes and virions due to their similar shape and size precluded quantification of the data. Therefore, we used immuno-gold labeling in which exosomes and SARS-CoV-2 virions were identified by an anti-CD63 antibody coupled to 18 nm gold particle and an anti-NP antibody coupled to a nanogold particle (1.4 nm), respectively (Fig 4B). We observed an enrichment of CD63 labeling around SARS-CoV-2 virions when incubated with ACE2+ exosomes, but not ACE2-exosomes (Fig. 4C). The pronounced clustering of CD63 in the ACE2+ exosome-virus mixtures may be indicative of multiple exosomes bound to the virions, however, we cannot definitively conclude the number of exosomes due to the lack of contrast of the exosome membranes.

**Figure 4.**
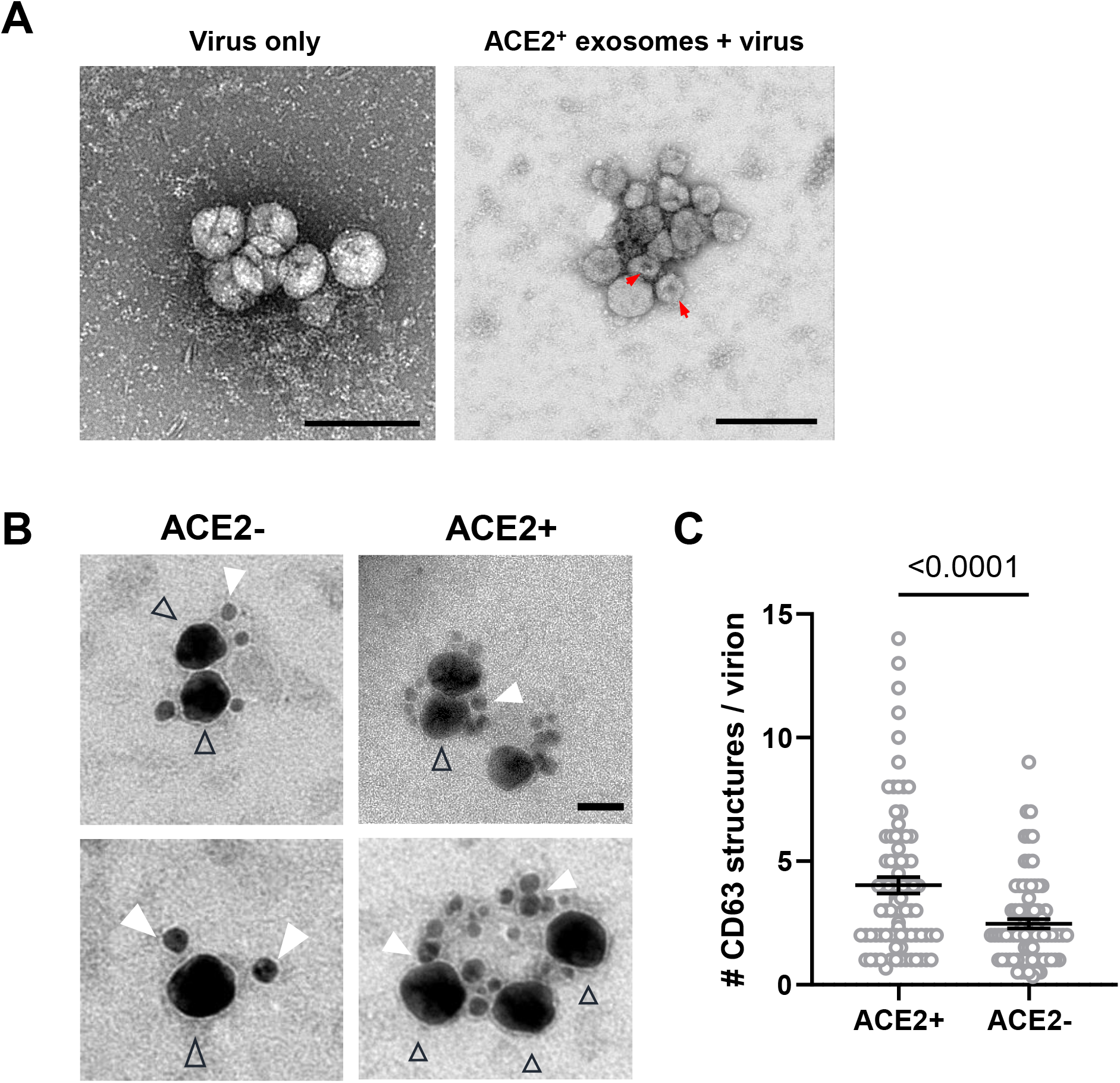
SARS-CoV-2 virions associate with ACE2+ exosomes. **(A)** Transmission electron microscopy (TEM) of negative stained exosomes and virus alone. Arrowheads: putative exosomes. Scale bars: 200nm. Red arrows mark the characteristic collapsing of exosomes during TEM imaging. **(B)** Double immunogold labeling TEM images of SARs-CoV-2 virions mixed with ACE2+ exosomes (right panel, scale bar: 50nm), or ACE2-exosomes (left panel). Virions are labeled using an antibody for SARS-CoV-2 N, followed by nanogold conjugated Fab (black triangle outline). Exosomes are labeled using an antibody against CD63, followed by 18nm colloidal gold conjugated IgG (filled white triangles). **(C)** Quantification of CD63 structures per virion present in a cluster containing both markers. ACE2+ (n=68 clusters), ACE2-(n=57). Mean ± SEM. Unpaired *t*-test with Welch’s correction. * *P* ≤ 0.05; ** *P* ≤ 0.01; *** *P* ≤ 0.001; **** *P* ≤ 0.0001. ns: not significant.

Lastly, we applied cryo-electron microscopy and tomography to confirm these associations between SARS-CoV-2 virions and ACE2+ exosomes. Virions were easily distinguished by the presence of protrusions that resembled the club-like head of the spike protein and exosomes were larger single membraned vesicles, spherical or ellipsoid in shape. Focusing on the area between the viral and exosome membranes, we were able to visualize an elongated structure resembling the post-fusion form of spike and ACE2 in some tomograms (Supp. Fig. 3A, 3B, Videos 1-2). Tomographic reconstruction also captured several virions around larger exosomes as well as potential virions inside an exosome (Video 3). Collectively, these data demonstrate that the presence of ACE2 on exosomes increases binding and clustering of virions.

## DISCUSSION

Our analyses of an inpatient cohort of COVID-19 patients revealed ACE2 levels on exosomes in BALF can display striking interindividual variability, and that a high proportion of ACE2+ exosomes, especially those with increased surface levels, are associated with reduced hospital stay. Although it is unclear whether ACE2+ exosomes are a marker or have a causative role in recovery from SARS-CoV-2 infection, our cell culture experiments indicate that they can inhibit viral infection by serving as defensosomes. The ability to produce sufficient numbers of defensosomes is key to providing optimal protection *in vitro*, and thus, understanding the factors that contribute to variable exosome production at mucosal surfaces during SARS-CoV-2 infection and other conditions will be of great interest. Transcriptional regulation and cleavage by proteases such as ADAM10 and ADAM17 [35] may also explain heterogeneity in ACE2+ exosomes. Other unanswered questions include how ACE2+ exosome release affects the availability of ACE2 in the tissue and whether this impacts acute injury to the lung, another function of this critical enzyme involved in cardiovascular processes [36]. Also, we focused on surface protein interactions, but it is important to consider other ways SARS-CoV-2 infection can be impacted by exosomes, which have been shown to facilitate the transfer of viral and cellular components to target cells with either pro-viral or antiviral consequences depending on the pathogen [37-39].

Infection by SARS-CoV-2 and other coronaviruses leads to formation of structures resembling autophagosomes [40-42], double-membrane vesicles generated during autophagy that target intracellular material to the lysosome for degradation [43]. Several SARS-CoV-2 proteins inhibit formation of autophagosomes or prevent the acidification of lysosomes and endosomes, promoting viral egress from the cell [44-49]. Although viruses such as SARS-CoV-2 evade autophagy-mediated degradation during intracellular replication, our results suggest extracellular virions may be subject to neutralization by defensosomes generated through an autophagy-like process dependent on ATG16L1. In this context, it is notable that defensosomes are induced by innate immune activation of TLRs that require a signaling endosome [50, 51], which may coincide with the location from which exosomes are generated. The termination of TLR7 signaling is mediated by the packaging of TLR7 into exosomes via a ESCRT independent mechanism involving ALIX and the recruitment of syntenin-1 by UNC93B1 [52], providing precedence for coupling viral RNA recognition with extracellular vesicle trafficking. Cells that produce significant amounts of IFN in response to sensing of viral PAMPs through TLRs, such as plasmacytoid dendritic cells [53], could play an important role in initiating early protection by activating cells in distal tissues to release exosomes.

Several clinical trials are underway for exosome-based therapies against various cancers and kidney disease [54-56]. The advantage of strategies involving exosomes includes their scalability, stability in biofluids, and ability to be engineered to carry desired receptors and even deliver drugs. Our results support a broad innate antimicrobial function for defensosomes that includes defense against viral disease. An improved understanding of defensosome regulation may inform clinical applications.

## SUPPLEMENTAL MATERIALS

**Supplemental Figure 1.**
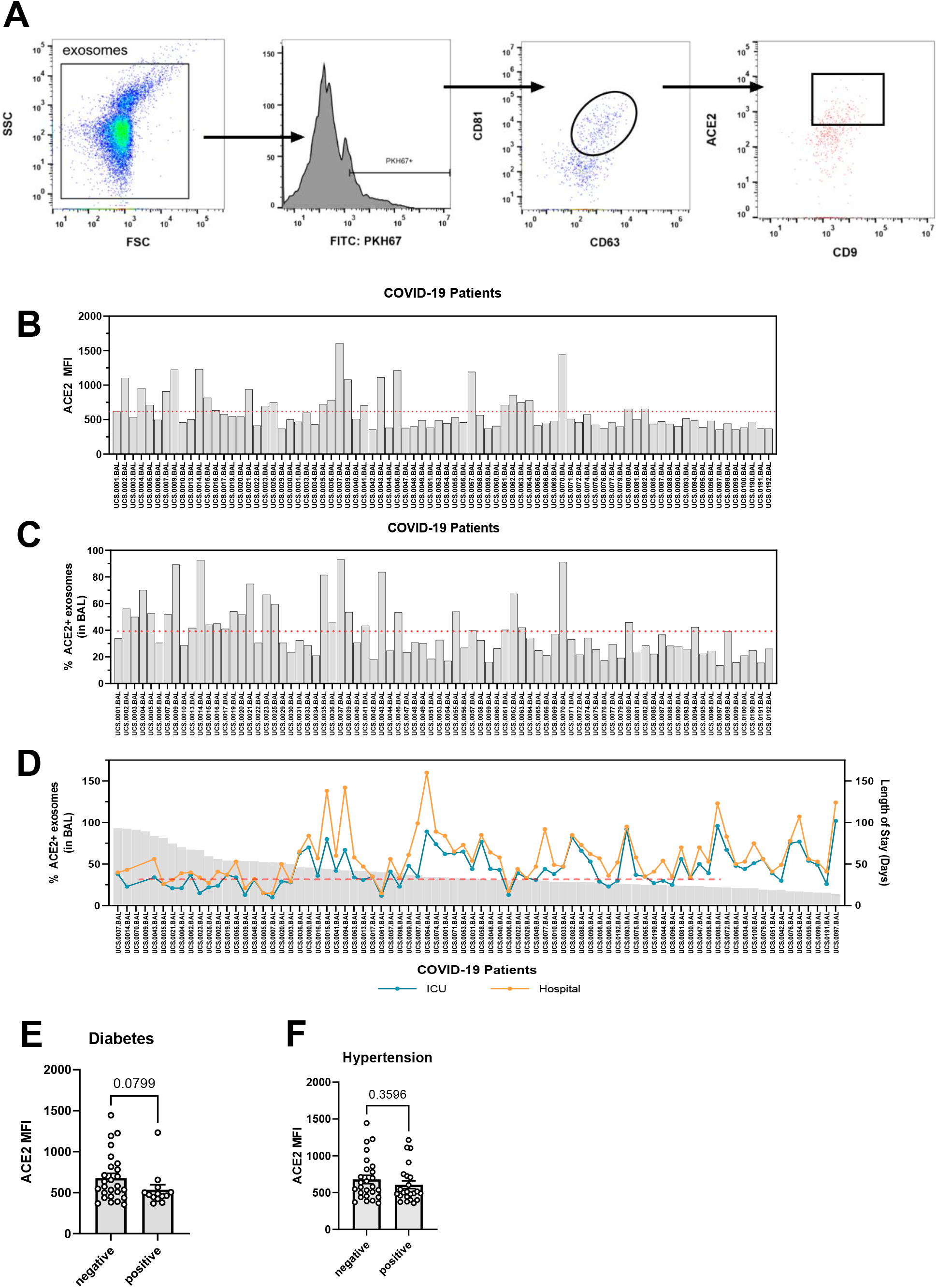
Characteristics of exosomes from COVID-19 patient BALF. **(A)** Gating strategy and representative flow cytometry plots from patient BALF. Exosomes were stained for antibodies against CD63, CD9, CD81, and ACE2. Exosomes were labeled with a lipid-intercalating dye, PKH67. FSC: forward scatter. SSC: side scatter. **(B)** Surface ACE2 on exosomes isolated from acellular BALF measured by mean fluorescence intensity (MFI) using flow cytometry. **(C)** Percentage of ACE2+ exosomes out of total exosomes in A. **(D)** Correlation between percentage of ACE2+ exosomes and length of stay in the ICU (orange line) and hospital (teal line) for all patients (including deaths) *(N=80)*. ACE2 levels on exosomes stratified by positive or negative patient status for **(E)** diabetes and **(F)** hypertension *(N=80)*. Red dotted lines indicate the average ACE2 MFI on ACE2+ exosomes in **B** or the average proportion of ACE2+ exosomes in **C** calculated from all COVID patients. * *P* ≤ 0.05; ** *P* ≤ 0.01; *** *P* ≤ 0.001; **** *P* ≤ 0.0001. ns: not significant.

**Supplemental Figure 2.**
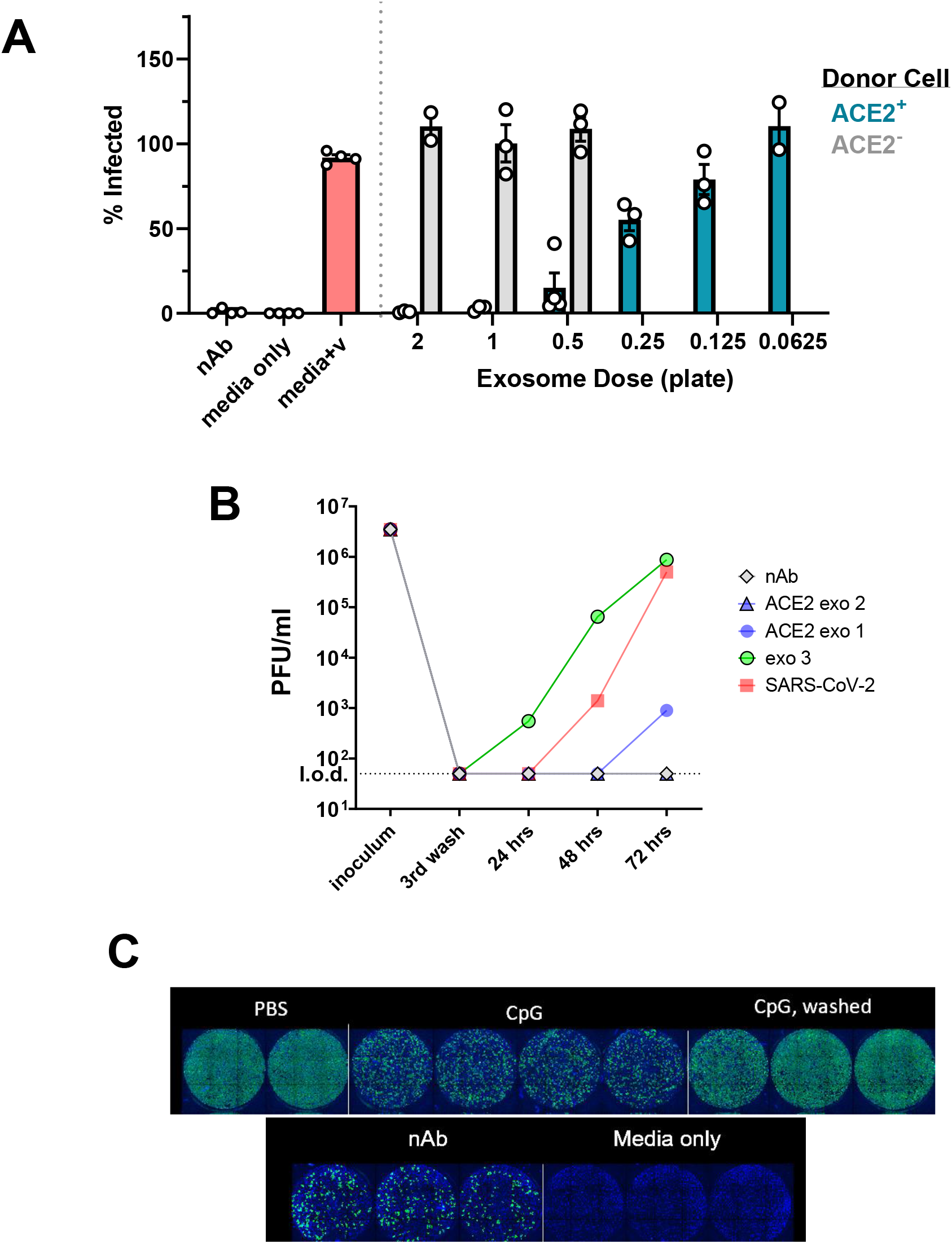
Characterizing role of exosomes against SARS-CoV-2 *in vitro*. **(A)** Infection level in Fig. 3D against number of donor A549^ACE2^ or A549 cells (in 15cm plates). **(B)** Number of infectious particles measured by plaque assay from apical washes every 24hr for 72hr of HAECs infected with SARS-CoV-2 alone (red), SARS-CoV-2 and ACE2+ exosomes (blue), SARS-CoV-2 and ACE2-exosomes (green), and SARS-CoV-2 with neutralizing Ab (grey). # corresponds to number of input plates (15cm). **(C)** Representative immunofluorescence images of Vero E6 cells infected with SARS-CoV-2 where the cells were stimulated/pretreated with PBS, CpG, or nAb from Fig. 3F.

**Supplemental Figure 3.**
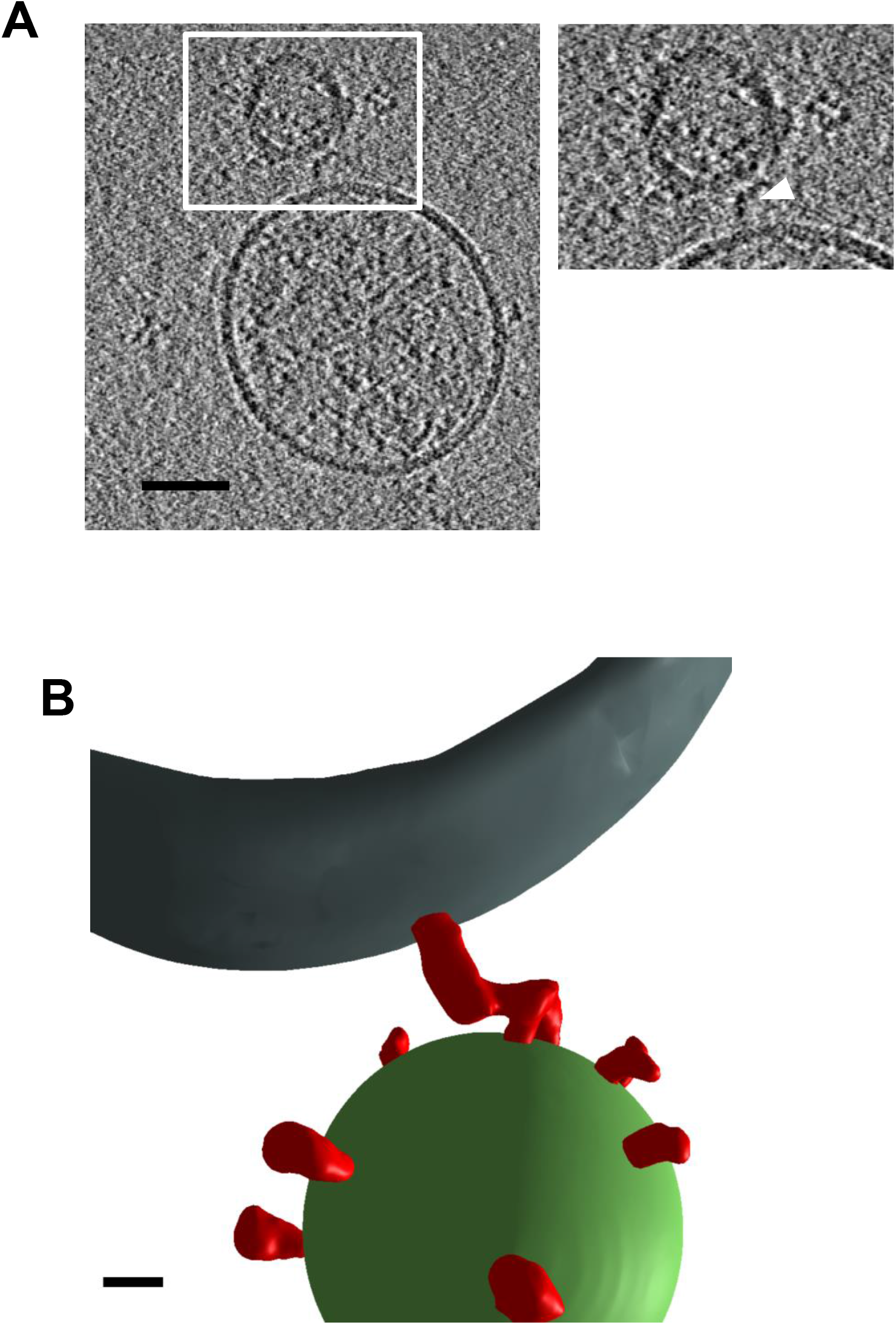
Cryo-EM tomogram and 3D rendering of an exosome and SARS-CoV-2 virion. **(A)** Representative tomographic slice of a SARS-CoV-2 virion and an exosome. Exosomes were isolated from ACE2+ A549 cells. Inset white arrow: spike, scale bar: 50nm. **(B)** Three-dimensional model of one individual exosome (gray) and SARS-CoV-2 virion with a membrane (green) and spike proteins (red) generated from segmentation. Scale bar: 10nm.

**Videos 1 and 2. Tomograms from Cryo-ET of SARS-CoV-2 and ACE2**^**+**^ **exosomes (1)** Scanning through all frames (forward and backward) of selected tomogram in Supp. 3A. Generated in ImageJ. **(2)** Rotational video of Supp. Fig. 4B. Each grid is a 10nm square. **(3)** Additional video of tomographic reconstructions of exosomes and virions.

## SUPPLEMENTARY TABLES

**Supplementary Table 1.** Demographics and clinical characteristics of the COVID-19 patient cohort.

| Variables | Cohort |
| --- | --- |
| **N** | 80 |
| **Age** | 64 [51-71] |
| **Age Category** |  |
| <51 | 19 (23.8%) |
| 51-60 | 11 (13.8%) |
| 61-70 | 27 (33.8%) |
| >70 | 23 (28.7%) |
| **Sex (Male)** | 62 (77.5%) |
| **Race** |  |
| White | 37 (46.2%) |
| Asian | 3 (3.8%) |
| Black or African American | 8 (10%) |
| Other | 32 (40%) |
| **Ethnicity** |  |
| Hispanic or Latino | 21 (26.2%) |
| **BMI** | 26.5 [24-30] |
| BMI $\geq 30$ | 24 (30%) |
| **Comorbidities** |  |
| Asthma | 1 (1.2%) |
| CAD | 12 (15%) |
| CHF | 6 (7.5%) |
| CKD | 11 (13.8%) |
| CVA | 15 (18.8%) |
| Diabetes | 31 (38.8%) |
| HLD | 26 (32.5%) |
| HTN | 41 (51.2%) |
| **Smoking Status** |  |
| Never | 53 (66.2%) |
| Ex.Smoker | 12 (15%) |
| Current | 6 (7.5%) |
| **Respiratory Culture** |  |
| Any respiratory pathogenic bacteria | 66 (82.5%) |
| *Staphylococcus aureus* | 15 (18.8%) |
| *C. albicans* | 17 (21.3%) |
| **Blood Culture** |  |
| Any positive bacterial culture | 46 (57.5%) |

| **Treatment and Outcomes** |  |
| --- | --- |
| Intubated | 80 (100%) |
| Dialysis | 20 (25%) |
| Dialysis days | 20 [9-32] |
| ECMO | 13 (16.2%) |
| ECMO days | 14 [8-71] |
| Ventilator days | 33 [24-49] |
| ICU length of stay | 39 [30-62] |
| Hospital length of stay | 53 [39-73] |
| **Mortality** | 19 (23.8%) |

Summary of hospitalized COVID-19 patients from which the ACE2+ exosomes were measured from the BALF. Data expressed as n(%) or Median[IQR]. BMI, body mass index. BMI is the weight in kilograms divided by the square of the height in meters. CHF: congestive heart failure; CAD: coronary artery disease; CKD: chronic kidney disease; CVA: cerebrovascular accident. HLD: hyperlipidemia; HTN: Hypertension; ECMO: extracorporeal membrane oxygenation. Respiratory culture: defined as having any respiratory culture performed. Positive bacteria: a culture resulting in any bacterial growth. Blood culture: defined as having any blood culture performed. ICU: intensive care unit; Ventilator days: total number of days on mechanical ventilation.

**Supplementary Table 2.** Regression using negative binomial model and length of stay in the ICU as the outcome.

| Length of stay in ICU (Days) |  |  |  |
| --- | --- | --- | --- |
| *Predictors* | *Incidence Rate Ratios* | *CI* | *p* |
| Age (years) | 1.0072 | 0.9907 – 1.0240 | 0.395 |
| % ACE2 positive (BAL exosomes) | 0.9865 | 0.9816 – 0.9915 | **<0.001** |
| Sex [M] | 2.5665 | 0.8587 – 7.6710 | 0.092 |
| Hypertension | 1.9773 | 0.2710 – 14.4269 | 0.501 |
| BAL *C. albicans* [Positive] | 1.5818 | 1.1825 – 2.1159 | **0.002** |
| Blood culture final result [Positive] | 1.1724 | 0.9698 – 1.4173 | 0.1 |
| **(Intercept)** | 46.6348 | 17.3749 – 125.1689 | <0.001 |
| **Observations** | 78 |  |  |
| **R<sup>2</sup> Nagelkerke** | 0.581 |  |  |

**Supplementary Table 3.** Linear regression on covariates including ACE2 MFI using length of stay in the ICU as the outcome.

| Length of Stay in ICU (Days) |  |  |  |
| --- | --- | --- | --- |
| *Predictors* | *Estimates* | *CI* | *p* |
| Age (years) | 0.4685 | -0.3995 – 1.3365 | 0.285 |
| ACE2 MFI | -0.0291 | -0.0481 – -0.0100 | **0.003** |
| sex [M] | 53.6830 | -3.6455 – 111.0116 | 0.066 |
| Hypertension | 22.1203 | -80.3790 – 124.6196 | 0.668 |
| BAL *C. albicans* [Positive] | 21.1636 | 5.5735 – 36.7537 | **0.009** |
| Blood culture final result [Positive] | 4.5791 | -5.4529 – 14.6112 | 0.366 |
| **(Intercept)** | 36.1704 | -14.3409 – 86.6817 | 0.158 |
| **Observations** | 78 |  |  |
| **R<sup>2</sup> / R<sup>2</sup> adjusted** | 0.273 / 0.165 |  |  |

**Supplementary Table 4.**
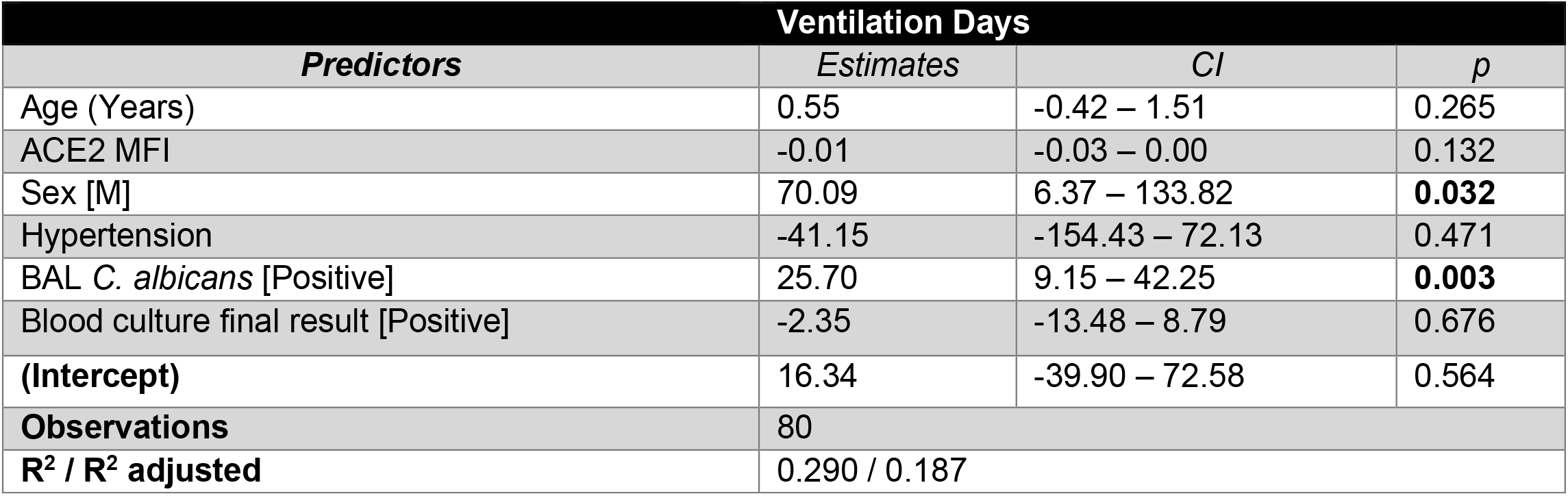
Linear regression on covariates including ACE2 MFI using ventilation days as an outcome.

| Ventilation Days |  |  |  |
| --- | --- | --- | --- |
| *Predictors* | *Estimates* | *CI* | *p* |
| Age (Years) | 0.55 | -0.42 – 1.51 | 0.265 |
| ACE2 MFI | -0.01 | -0.03 – 0.00 | 0.132 |
| Sex [M] | 70.09 | 6.37 – 133.82 | **0.032** |
| Hypertension | -41.15 | -154.43 – 72.13 | 0.471 |
| BAL *C. albicans* [Positive] | 25.70 | 9.15 – 42.25 | **0.003** |
| Blood culture final result [Positive] | -2.35 | -13.48 – 8.79 | 0.676 |
| **(Intercept)** | 16.34 | -39.90 – 72.58 | 0.564 |
| **Observations** | 80 |  |  |
| **R<sup>2</sup> / R<sup>2</sup> adjusted** | 0.290 / 0.187 |  |  |

**Supplementary Table 5.** Regression using negative binomial model and ventilation days as the outcome.

| Ventilation Days |  |  |  |
| --- | --- | --- | --- |
| *Predictors* | *Incidence Rate Ratios* | *CI* | *p* |
| Age (Years) | 1.0153 | 0.9947 – 1.0363 | 0.146 |
| % ACE2 positive (BAL exosomes) | 0.9923 | 0.9869 – 0.9979 | **0.007** |
| Sex [M] | 5.3818 | 1.4056 – 20.6056 | **0.014** |
| Hypertension | 0.4474 | 0.0416 – 4.8157 | 0.507 |
| BAL *C. albicans* [Positive] | 1.6952 | 1.2044 – 2.3859 | **0.002** |
| Blood culture final result [Positive] | 1.0380 | 0.8232 – 1.3089 | 0.752 |
| **(Intercept)** | 20.4347 | 6.0498 – 69.0235 | **<0.001** |
| **Observations** | 80 |  |  |
| **R<sup>2</sup> Nagelkerke** | 0.483 |  |  |

## MATERIALS AND METHODS

### Cohort of critically ill COVID-19 patients

The characteristics of this cohort have recently been described [22]. Briefly, we enrolled adults (>18 years old) admitted to the intensive care unit due to respiratory failure requiring invasive mechanical ventilation with confirmed diagnosis of SARS-CoV-2 infection by reverse transcriptase polymerase chain reaction (RT-PCR). Lower airway samples (bronchoalveolar lavage, BAL) were obtained during clinically indicated bronchoscopy performed for airway clearance or for percutaneous tracheostomy placement. Surviving subjects signed informed consent to participate in this study while samples and metadata from subjects who died or were incapacitated were de-identified and included this study. All patients or their legal representative agreed to participate via our NYU IRB approved protocol (IRB no. s16-00122/01598). The study protocol was approved by the Institutional Review Board of New York University.

### Cell Culture and lentivirus production

A549 cells were purchased from ATCC. A549^ACE2^ cells were generated using lentiviral transduction of a human ACE2 cDNA expressing plasmid (backbone: pLV-EF1a-IRES-Puro (Addgene 85132) as previously described [31]. TLR7+ A549 were generated by lentiviral transduction of pcDNA3-TLR7-YFP (a gift from Doug Golenbock (Addgene plasmid # 13022; http://n2t.net/addgene:13022; RRID:Addgene_13022)). A549^ACE2^ cells were selected using 1μg/ml puromycin. Cells were cultured with 10% Dulbecco’s modified Eagle medium (DMEM (Gibco), 10% fetal calf serum (FCS), 1% non-essential amino acids (NEAA), 1% Penicillin/Streptomycin) and filtered using a 0.22μm filter unit (Millipore). Vero E6 cells were purchased from ATCC (cat. No. CLR-1586) and maintained in same media as A549 cells, but including 10mg/mL Amphotericin B. 293FTs were maintained in same media as A549 cells. Lentivirus was produced by transfecting 10cm plate of 293FTs with 10μg of plasmid using Lipofectamine 3000. Media was exchanged the next day and lentivirus was collected the following day. 1mL lentivirus was used to transduce A549 cells using Polybrene (Millipore). TKO (IFNAR1/IFNLR/IFNGR1) A549 cells were generously provided by Marco Binder at the German Cancer Research Center (DKFZ), Heidelberg, Germany [57].

Stimulations were done with 0.5μM CpG-ODN 2216 (Class A) or CpG-ODN 2006 (Class B) (Invivogen), 1μg/mL R848 (Invivogen), human IFN-β (R&D), IFN-λ2 (cat. No. 8417-IL-025, R&D), IFNα (cat. No 11200-2, R&D), 1μg/mL 3’3’-cGAMP, 1ug/mL 2’3’-cGAMP (Invivogen), and PolyI:C LMW and HMW with and without LyoVec (Invivogen).

### BSL3

Plaque and neutralization assays were performed in the BSL3 facility of NYU Grossman School of Medicine, in accordance with its Biosafety Manual and Standard Operating Procedures.

### Plaque Assay

Viral titers were determined by plaque assay. Vero E6 cells were seeded into 12-well plates to reach confluency the next day. When confluent, cells were washed twice with PBS (Ca/Mg+) before the infection with 1:10 serial dilutions of the samples for 1 h at 37°C. At 1 h post virus addition, virus was removed, cells were washed again twice with PBS (Ca/Mg+), and overlay-medium was added (minimal essential medium (MEM), 1.5µg/ml Trypsin-TPCK, 1.6% Oxoid-Agar, 1% Pen-Strep, 1% GlutaMax, 20 mM HEPES, 0.4% BSA, and 0.24% NaHCO3). At 72 h post infection, the cells were fixed by adding 10% formalin solution for 24 hrs. After fixation, overlay was removed and cells were stained with crystal violet solution (0.1 % crystal violet in 1.9% Ethanol and 19% Methanol in cell culture grade H2O) for at least 30 min. Plates were rinsed once with H2O before determining the viral titer.

### Virus Amplification

All SARS-CoV-2 stock preparations and subsequent infection assays were performed in the CDC/USDA-approved biosafety level 3 (BSL-3) facility in compliance with NYU Grossman School of Medicine guidelines for BSL-3. SARS-CoV-2 isolate USA-WA1/2020, deposited by the Centers for Disease Control and Prevention, was obtained through BEI Resources, NIAID, NIH (cat. no. NR-52281; GenBank accession number MT233526). The USA-WA1/2020 stock was passaged twice in Vero E6 cells to generate a passage 6 working stock (1.0E+06 PFU/ml) for our studies.

### *In vitro* Neutralization Assay

3.2×10^4^ Vero E6 cells were seeded in black 96-flat well with clear bottom plates 24hr before infection. SARS-CoV2 was mixed 1:1 v/v with exosomes in infection media (DMEM, 10% HEPES, 10% NEAA) and allowed to bind for 1hr at 37°C. Heat-inactivated plasma (56°C for 1 hour) from a convalescent patient was used as a positive control. Confluent Vero cells were then infected with this exosome-virus mixture. At 24 hpi, plates were fixed with 10% formalin solution for 30-45 min and then washed with dH2O. Plates were permeabilized with 0.1% Triton-X 100, blocked with 1% BSA/PBS and stained for SARS-CoV-2 N mouse monoclonal SARS-CoV-2 anti-N antibody 1C7 (1:1000, kind gift of Thomas Moran) overnight at 4°C. Plates were then incubated with Alexa Fluor 647 goat anti-mouse Ab (1:2000) and DAPI (1:2000) at 37°C for 1hr. After 3 washes with PBS, plates were imaged using the Cell Insight CX7 LZR high-content screening platform. Images were analyzed and quantified with HCS Navigator software.

### Infection and immunofluorescence of HAECs

Maintenance of HAECs were done as described in [31]. At 48 h prior to infection, 6-8-week-old HAECs were washed apically twice for 30 min each with prewarmed PBS (Ca/Mg+). At 1 hr prior to infection, cultures were washed apically twice for 30 min each with prewarmed PBS. Each culture was infected with a viral dilution containing with 1.35E+05 PFU with or without exosomes for 2 h at 37°C. The inoculum was removed, and the cultures were washed three times with prewarmed PBS. For each washing step, PBS was added to the apical surface, and cultures were incubated at 37°C for 30 min before the PBS was removed. The third wash was collected and stored at −80°C for titration by plaque assay on Vero E6 cells. Infectious particles were collected every 24 hr (up to 72 hr) by adding 60 μl of prewarmed PBS, incubating at 37°C for 30 min, and washes were stored at −80°C. Wells were fixed by submerging in 10% formalin for 24 h and washed three times with PBS before analysis by immunofluorescence. Fixed HAECs were quenched 50 mM NH_4_Cl (in PBS), permeabilized using a solution of, 0.1% Saponin, and blocked with 2% BSA. Cultures were stained with anti-SARS nucleocapsid protein antibody which cross reacts with SARS-CoV-2 N (1:1000, cat. no. 200-401-A50; Rockland) and goat-anti-rabbit Alexa Fluor 488 and DAPI.

### Mitochondrial DNA generation

Short mitochondrial DNA fragments (221bp) were amplified from gDNA from A549 cells with and without 8-Oxo-2’-deoxyguanosine-5’-Triphosphate (TriLink Biotechnologies) using GoTaq Mastermix. Forward primer: 5’-CCCCACAAACCCCATTACTAAACCCA-3’. Reverse: 5’-TTTCATCATGCGGAGATGTTGGATGG-3’. Fragments were isolated using Quick PCR Clean Up kit (Qiagen). Genomic DNA fragments were generated by sonicating genomic DNA isolated from A549 cells (Qiagen Blood and Tissue DNeasy kit) using genomic DNA tips (20g).

### shRNA knockdown

Lentivirus based knockdown of human *ATG16L1* (*5’-*ACTGTAGCTTTGCCGTGAATG-*3’*) was performed using MISSION shRNA constructs (Sigma-Aldrich) and psPAX2 packaging system. psPAX2 was a gift from Didier Trono (Addgene plasmid # 12260). Lentivirus expressing shRNAs were produced by DNA transfection using Lipofectamine 3000 (Thermo Fisher). Transduction was performed using Polybrene (Millipore) and selection was done using 1μg/mL puromycin.

### Flow Cytometry

Exosomes pellets were stained with 100μl of an antibody cocktail in 1X PBS (Corning) containing anti-CD9 (human HI9a, mouse MZ3), anti-CD63 (human H5C6, mouse NVG-2), anti-CD81 antibodies (human 5a6, mouse Eat-2) from Biolegend, and anti-human ACE2 (R&D Biosystems cat no. FABAF9332R) at 1:100 for 1hr at 4°C, rocking. Exosomes were stained with PKH67 cell linker dye (Sigma). Exosomes were washed with 1X PBS once and ultracentrifuged at 100,000x*g* for 1.5hr. The exosome pellet was resuspended in PBS. Cells and exosomes were analyzed using a Beckman Coulter Cytoflex S cytometer at the slowest flow rate.

### Exosome Isolation

Cell culture supernatants were spun at 500x*g* for 10min, followed by a 10,000x*g* spin for 10min. Supernatants were passed through a 0.22μm filter (Millex-GP PES). Supernatants were washed with 1x PBS and pelleted by ultracentrifugation at 100,000x*g* for 1.5hrs. Exosomes used for neutralization were resuspended in viral media (10% DMEM, 1% HEPES, 1% PenStrep) and transferred to a non-TC round bottom 96-well plate (Corning).

All human BAL samples were processed into cellular and supernatant fractions before freezing for further analysis. Exosomes were isolated from 200μl of thawed acellular BAL fluid and fixed to a final concentration of 4% PFA before being removed from the BSL-3 facility. Samples were washed with PBS and ultracentrifuged at 100,000xg for 1.5hr before staining for flow cytometry analysis.

### Negative Stain, Immunogold Labeling, and Cryo Electron Tomography

Exosomes isolated from A549 and A549^ACE2^ cells stimulated with Bafilomycin A1 were mixed with SARS-CoV-2 and allowed to bind for 1hr at 37°C in the BSL3 before being fixed with 4% PFA. Suspensions of SARS-CoV-2/exosomes were ultracentrifuged once at 100,000 x *g* to pellet, and then resuspended in PBS before processing for TEM imaging.

Negative stain of exosome-virus mixtures involved deposit of 3μl of 4% paraformaldehyde fixed exosomes on glow discharged carbon copper grids. Grids were stained with 1% uranyl acetate and were imaged under FEI Talos L120C TEM and photographed with a Gatan (4*k* × 4*k*) OneView camera (Gatan Inc., Pleasanton, CA.)

For double immunogold labeling, exosome-virus mixtures in PBS were fixed to final concentration of 4% PFA and placed on glow-discharged formvar-carbon copper grids for 20min. Staining was done as follows: Grids were washed with PBS (pH 7.4) twice for 2 min each followed by incubation with PBS/Glycine 50mM for 3 min. Grids were permeabilized and blocked with 0.1% Saponin/1% cold-water fish skin gelatin (Perm buffer) for 10 min and first stained with anti-human CD63 (Abcam, ab59479) antibody (1:10) in Perm buffer for 1.5hr. Grids were washed 6x with PBS with 0.1% Saponin for 2 min, and then incubated with 18nm gold conjugated anti-mouse antibody (18nm colloidal gold AffiniPure goat anti-mouse IgG (H+L), Jackson ImmunoResearch Laboratories, Inc., West Grove, PA), in Perm buffer for 30min. After washing with PBS, grids were fixed in 2% PFA in PBS for 5 min.

Grids were washed with PBS and then incubated in PBS/50mM glycine for 3min to quench the unbounded aldehyde group. The grids were incubated with anti-NP antibody (1:100) in perm buffer for 1.5 hr followed by anti-mouse Fab’ nanogold (1:250, Nanoprobe, New York) in perm buffer for 30min. Grids were fixed in 1% glutaraldehyde in PBS for 5min, washed with distilled water, followed by washes with 0.02M sodium citrate buffer (pH 7.0). The silver enhancement is done using HQ Silver enhancement kit (#2012, HQ Silver, Nanoprobe, New York) for 9 minutes, and rinsed with ddH2O. Finally, grids were contrasted with 3% uranyl acetate and embedded with 2% methylcellulose in a ratio of 1 to 9.

Immunogold and nanogold double labeling was quantified using ImageJ software (NIH).

### CryoEM Sample Collection

Samples for CryoET were plunge frozen in liquid ethane with a VitroBot Mark IV (Thermo Fisher) at 20°C at 100% humidity on C-flat (Protochips, Morrisville NC) 1.2/1.3 grids with 80s easyGlow (Ted Pella Inc, Redding CA) glow discharge. All images were acquired on the Talos Arctica (Thermo Fisher) housed at NYU. Images were taken at 200 kV with a Gatan K3 imaging system (Gatan Inc, Pleasanton CA) collected at 28,000X nominal magnification. The calibrated super-resolution pixel size of 0.7040 Å was used for processing.

Micrographs were collected using Leginon [58, 59] at a dose rate of 11.21 e^-^/Å^2^/s with a total exposure of 3.50 seconds, for an accumulated dose of 39.23 e^-^/Å^2^. Intermediate frames were recorded every 0.07 seconds for a total of 50 frames per micrograph. Frames were aligned using Motioncor2 [60].

Tilt series were collected using SerialEM [61] at a dose rate of 11.21 e^-^/Å^2^/s with a total exposure of 0.25 second for an accumulated dose of 2.79 e^-^/Å^2^ per tilted image. Intermediate frames were recorded every 0.05 seconds for a total of 5 frames per micrograph. Movies were collected in super-resolution mode. Tilt series were acquired in a bidirectional pattern starting from −18°, with a range from −60° to 60° in 3° step with a total dose of 114e^-^/Å^2^.

### Tilt series processing

Frames were aligned using Warp [62], binned to the physical pixel size of 1.408 Å, and constructed into a tilt series stack for further processing. Micrographs within each tilt series were aligned with EMAN2 [63] and constructed into a tomogram at binning 4 (5.63 Å/pixel) using Fourier inversion. Tomograms were denoised using Topaz [64]. Denoised tomograms were inspected using IMOD [65] and clipped to appropriate size. Movies were made from selected tomograms using ImageJ. Automated segmentation was performed using EMAN2 to segment membranes, and further 3D rendering and refinement of Spike proteins was done manually on ORS Dragonfly (Object Research Systems, Quebec, Canada).

### Patient correlation analysis

Anonymized data contained patient demographics, medications prescribed during hospital admission, vitals upon entry to the hospital, reported comorbidities, spike and RBD antibody titers, and blood and lung culture results. All analyses were performed in R Studio (version 1.4.1717).

### Statistical Analyses

Statistical analyses were performed using GraphPad Prism 9.0 (GraphPad Software, San Diego, California USA; https://www.graphpad.com/). All graphs show the mean and standard error of the mean (SEM). An unpaired two-tailed t-test was used to evaluate differences between two groups with Welch’s correction. One way ANOVA with Dunnett’s post-test was used to evaluate experiments involving groups of 3 or more. Two-way ANOVA with Dunnett’s or Šidák’s post-test was used to evaluate groups with more than one variable. Correlation was analyzed using Pearson r. For each test, a p-value <0.05 is considered statistically significant. Clinical data was analyzed using R-studio. We performed descriptive analysis extracting data from EHR for these patients and summarizing this data with R. Linear model was used with the outcome as length of stay in the ICU. As a sensitivity analysis, we also performed a negative binomial model. with ACE2 MFI as a predictor instead of % ACE2 positivity, which yielded the same results.

## ACKNOWLEDGEMENTS

We thank the Office of Science & Research High-Containment Laboratories at NYU Grossman School of Medicine for their support in the completion of this research and the NYU Langone Health’s Microscopy Laboratory and CryoEM Core. We also thank Marco Binder’s group for providing the TKO A549 cells, Rachel Prescott for maintaining HAECs, and Stephen T. Yeung and Yonghua Li for thawing and handling BALF samples. Supplemental figure 4A was created using BioRender.

## AUTHOR CONTRIBUTIONS

K.L.C., M.D., V.J.T., and K.C. conceived and designed the study. K.L.C., M.D.V., J.G., C.B. performed the experiments and analyzed the data. K.L.C., K.D., and J.S., performed and analyzed the electron microscopy data. F.-X.L provided assistance TEM and W.J.R. performed and analyzed CryoEM experiments. L.N.S provided BAL samples and patient metadata, L.N.S. and B.S. provided guidance on analyzing patient data. L.E.T. provided advice on statistical analyses of patient metadata. K.L.C., J.G. M.D.V., M.D., V.J.T., and K.C. interpreted the data. K.L.C., V.J.T., and K.C. wrote the paper with input from all authors.

